# Amino acid transporter B^0^AT1 influence on ADAM17 interactions with SARS-CoV-2 receptor ACE2 putatively expressed in intestine, kidney, and cardiomyocytes

**DOI:** 10.1101/2020.10.30.361873

**Authors:** Jacob T. Andring, Robert McKenna, Bruce R. Stevens

## Abstract

SARS-CoV-2 exhibits significant experimental and clinical gastrointestinal, renal, and cardiac muscle tropisms responsible for local tissue-specific and systemic pathophysiology capriciously occurring in about half of COVID-19 patients. The underlying COVID-19 mechanisms engaged by these extra-pulmonary organ systems are largely unknown. We approached this knowledge gap by recognizing that neutral amino acid transporter B^0^AT1 (alternately called NBB, B, B^0^ in the literature) is a common denominator expressed nearly exclusively by three particular cell types: intestinal epithelia, renal proximal tubule epithelium, and cardiomyocytes. B^0^AT1 provides uptake of glutamine and tryptophan. The gut is the main depot expressing over 90% of the body’s entire pool of SARS-CoV-2 receptor angiotensin converting enzyme-2 (ACE2) and B^0^AT1. Recent cryo-EM studies established that ACE2 forms a thermodynamically favored dimer-of-heterodimers complex with B^0^AT1 assembled in the form of a dimer of two ACE2:B^0^AT1 heterodimers anchored in plasma membranes. Prior epithelial cell studies demonstrated ACE2 chaperone trafficking of B^0^AT1. This contrasts with monomeric expression of ACE2 in lung pneumocytes, in which B^0^AT1 is undetectable. The cell types in question also express a disintegrin and metalloproteinase-17 (ADAM17) known to cleave and shed the ectodomain of monomeric ACE2 from the cell surface, thereby relinquishing protection against unchecked renin-angiotensin-system (RAS) events of COVID-19. The present study employed molecular docking modeling to examine the interplaying assemblage of ACE2, ADAM17 and B^0^AT1. We report that in the monomer form of ACE2, neck region residues R652-N718 provide unimpeded access to ADAM17 active site pocket, but notably R708 and S709 remained >10-15 Å distant. In contrast, interference of ADAM17 docking to ACE2 in a dimer-of-heterodimers arrangement was directly correlated with the presence of a neighboring B^0^AT1 subunit complexed to the partnering ACE2 subunit of the 2ACE2:2B^0^AT1] dimer of heterodimers, representing the expression pattern putatively exclusive to intestinal, renal and cardiomyocyte cell types. The monomer and dimer-of-heterodimers docking models were not influenced by the presence of SARS-CoV-2 receptor binding domain (RBD) complexed to ACE2. The results collectively provide the underpinnings for understanding the role of B^0^AT1 involvement in COVID-19 and the role of ADAM17 steering ACE2 events in intestinal and renal epithelial cells and cardiomyocytes, with implications useful for consideration in pandemic public hygiene policy and drug development.

## INTRODUCTION

While lung pathophysiology is the hallmark of COVID-19, the SARS-CoV-2 virus exhibits significant experimental and clinical gastrointestinal (GI), renal, and cardiac muscle tropisms that invoke specific local and systemic inflammasome pathophysiology [1–7]. These extra-pulmonary engagements occur capriciously in about half of COVID-19 cases, often appearing before fever onset, and extending after pulmonary symptoms abate [1–13]. The underlying mechanisms of COVID-19 involving these three cell types is largely unknown.

We approached this knowledge gap by recognizing that the amino acid transporter B^0^AT1, serving such substrates as glutamine and tryptophan, is a common denominator uniquely expressed in the plasma membranes of epithelial cells of intestine and renal proximal tubule and in cardiomyocytes, but is undetectable in lung pneumocytes [1–7, 10, 14–36]. The human GI tract is the body’s site of greatest magnitude expression of both B^0^AT1 and the angiotensin converting enzyme-2 (ACE2) receptor for SARS-CoV-2, based on expression patterns from single cell RNA seq, immunohistochemistry, and functional genomics studies [10, 14–19, 37]. These cell types also express a disintegrin and metalloproteinase-17 (ADAM17), a key player in the landscape of COVID-19 pathophysiology [1, 2]. The present study addresses the interplaying triangulation of these three proteins.

B^0^AT1 was originally discovered and functionally characterized by Stevens and coworkers [20–27] as the Na^+^-coupled neutral amino acid transport system in intestinal epithelial apical brush border membranes (alternately called NBB, B, B^0^ in the literature [21]), then subsequently Ganaphathy, Broer, Fairweather, Verrey, and colleagues [28–34, 38] cloned its SLC6A19 gene leading to experimental evidence of trafficking/chaperoning of B^0^AT1 by ACE2 to the cell surface. Building from these studies, Yan and coworkers in Zhou’s lab [35] utilized cryoelectron microscopy to determine a 2.9 Å complex of B^0^AT1 with ACE2, and established that in the presence of B^0^AT1, ACE2 forms a thermodynamically favored dimer-of-heterodimers complex assembled as two ACE2:B^0^AT1 heterodimers anchored in plasma membranes.

In contrast to B^0^AT1-expressing cells, lung pneumocyte membranes lack B^0^AT1 and express post-translationally mature ACE2 as a stand-alone monomer [1, 2]. Regardless of the organ—whether in lung, cardiopulmonary, renal or gastrointestinal tissues—the ACE2 membrane-tethered ectodomain possesses peptidyl carboxypeptidase activity that hydrolyzes nutritive and bioactive peptides, notably angiotensin *II* (Ang*II*), thus serving as protection against overactivity of the pernicious arm of the renin-angiotensin-system (RAS) [1–7, 10, 12, 14–36, 39–52]. SARS-CoV-2 hijacks the membrane-attached ACE2 ectodomain as its receptor and staging arena for priming the spike S-protein leading to virion cell entry [1, 2].

ADAM17 is a membrane-anchored enzyme that is central to mediating a variety of proinflammatory and signaling events in epithelial and endothelial cell types. ADAM17 possesses a metzincin-like HELGHNFGAEHD active site motif that hydrolyzes nearly 100 different substrates [53]. This serves a “sheddase” function that proteolytically releases the ectodomains of membrane-bound proteins into the extracellular milieu, with the consequence that ADAM17 activity leads to pathophysiological downsides in cardiopulmonary, GI, and renal systems [54].

ADAM17 cleaves monomeric ACE2 [47, 49, 55], resulting in separation and release of the soluble ACE2 (sACE2) ectodomain from the membrane anchor stalk, thereby relinquishing its protective roles against RAS. ADAM17 is activated by Ang*II*, and therefore SARS-CoV-2 induces an ADAM17-triggered positive feedback viscous cycle escalating pernicious Ang*II* levels unabated by ACE2 which is progressively inhibited [12]. These events of monomeric ACE2 contribute to the morbidity and mortality of COVID-19 [1, 2].

There is no consensus in the literature regarding the specific ACE2 residues targeted by ADAM17 involved in release of the ectodomain. Putative cleavage site bonds and surface residue attraction targets have been posited to lie within a broadly-defined region spanning residues R652-I741 within the neck region [47, 49, 55], based on biochemical analyses of *in vitro* expression of monomeric ACE2. The role of B^0^AT1 in influencing the ADAM17-ACE2 relationship is unreported. To address the many important knowledge gaps in tissue-specific manifestations of COVID-19, the present projected employed molecular docking modeling to examine ADAM17’s active site pocket access to ACE2, and the role of B^0^AT1 influencing the ADAM17-ACE2 relationship.

## METHODS

Online protein-protein docking software ClusPro 2.0 [56–58] was employed for docking simulations, with ADAM17 (PDB ID:3LGP chain A) designated as the ligand for each paired interaction. The active site of the catalytic domain of ADAM17 is a member of the metzincins with the **HE**XX**H**XXGXX**H**D motif, such that ADAM17 specifically possess a zinc atom held by H405, H409, H415 in conjunction with glutamate at E406 serving an acid/base catalytic function[53]; therefore these four residues were chosen as ADAM17 docking “attraction” residues. Targets included PDB ID: 6M17 [35] employed as a 2ACE2:2B^0^AT1 dimer-of-heterodimers complexed with SARS-CoV-2 receptor binding domain (RBD), and PDB:6M18 [35] employed analogously lacking SARS-CoV-2 RBD. Additional targets included ACE2 monomers and 1ACE2:1B^0^AT1 heterodimers derived from these structures using PyMOL[59]. Prior to docking, the coordinate files were modified by removing small molecules including solvent, inhibitors, ligands, glycosylation molecules, diphosphoglycerate, and non-standard amino acids. For each docking simulation, non-physiologically relevant domains of the complexes were masked out using PyMOL [59] to prevent non-specific binding (for example, hydrophobic residues within the transmembrane domain of membrane anchors).

ClusPro docks protein pairs in a three-step system [56–58]: 1) rigid body docking placed through 70,000 rotations; 2) clustering of low energy complexes; 3) energy minimization. The docked complexes are scored based on ClusPro 2.0 equation of the general form,

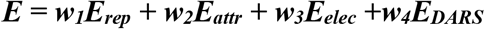

where *E_rep_* is energy of repulsion from van der Waals interactions, *E_attr_* is energy of attraction from van der Waals interactions, *Eelec* is the electrostatic energy term, and *E_DARS_* is the energy of desolvation. Docking scoring was based on the “Balanced” option choice with constants *w_1_* = 0.4, *w_2_* = −0.4, *w_3_* = 600, *and w_4_* = 1.0. ClusPro clusters together the top scored 1000 docked positions by finding a local center with the most docking neighbors within a 9Å sphere [58]. Energy minimization within a given cluster is used to remove any steric clashes between side chains, with the top docking scores outputted along with the corresponding complexed structure coordinate files [57, 58]. The best model chosen from among the output options resulting from each ClusPro docking was based on the highest ranked most populated cluster center (cluster size), rather than based on individual lowest energy structures, as recommended by Vajda and colleagues and CAPRI experiment data [56–58]. PDBePISA [60] was used to determine the binding contact interface residues. Figures were generated using PyMOL[59].

## RESULTS

The fundamental ACE2 monomer organization [1, 2] is shown in Fig. 1A, whereby a hydrophobic transmembrane stalk within the plasma membrane anchors and tethers the ACE2 ectodomain to the cell surface via a neck region. Molecular modeling results of the present study collectively implicated a span of ACE2 residues covering R652-N718 within the neck region (Fig. 1B) that provided the target for attempted dockings by ADAM17 as the ligand. In the case of monomeric ACE2, Fig. 2 shows the successful docking model, whereby ADAM17 catalytic zinc atom and its corresponding active site pocket residues H415 and E406 were attracted <2-3 Å to ACE2 residues R652, K657 and K659, but notably >10-15 Å distant to ACE R708 and S709.

**Fig. 1.**
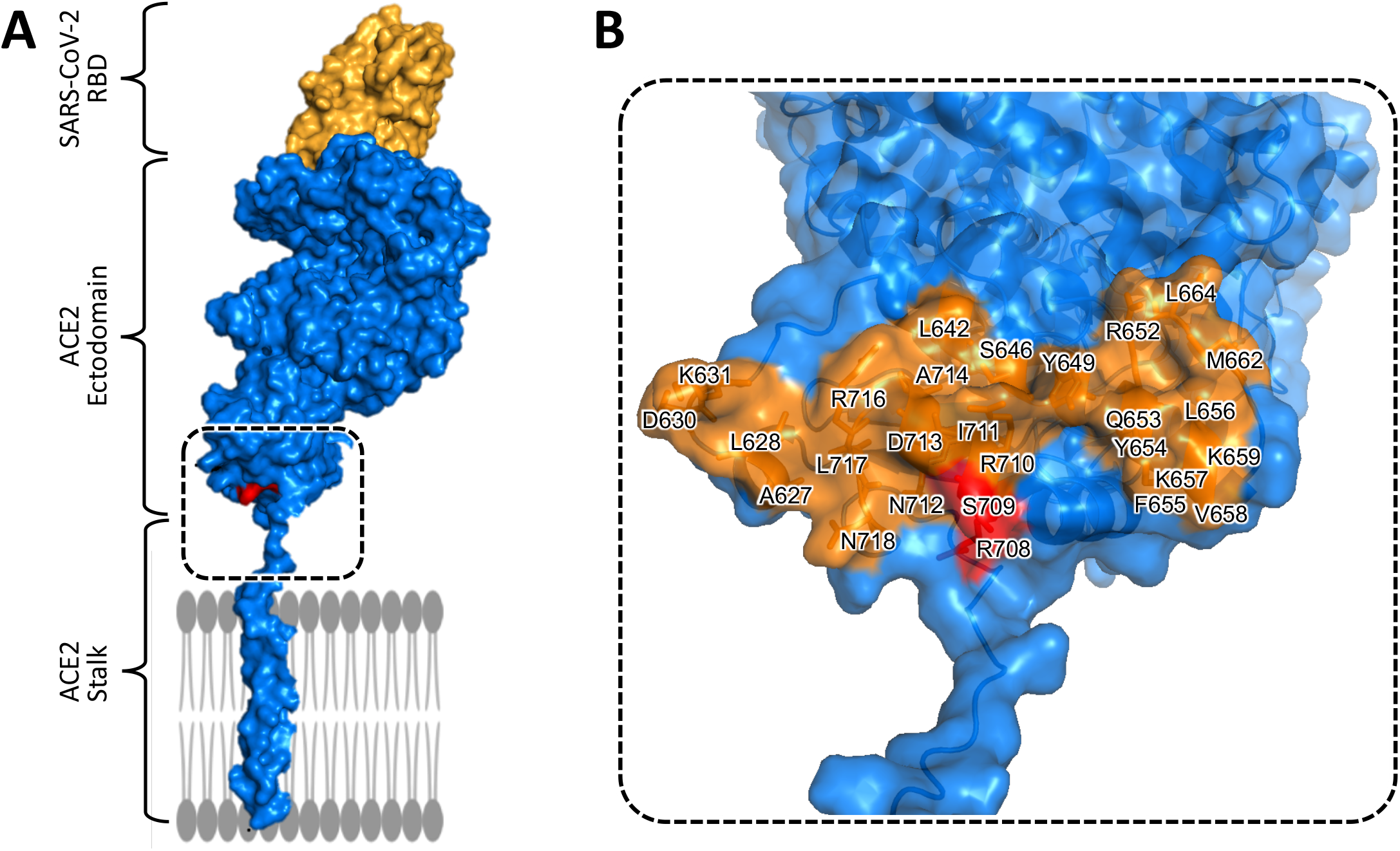
ACE2 monomer with neck region residues bridging ectodomain to anchor. **A.** ACE2 monomer (blue) shown with hydrophobic membrane anchoring stalk, and with the ectodomain complexed with SARS-CoV-2 receptor binding domain (RBD, gold). Box denotes ACE2 neck region spanning residues R652-N718. **B.** Closeup of box in (A) labeled with key neck region (brown) residues spanning R652-N718, which includes R708 and S709 residues (red).

**Fig. 2.**
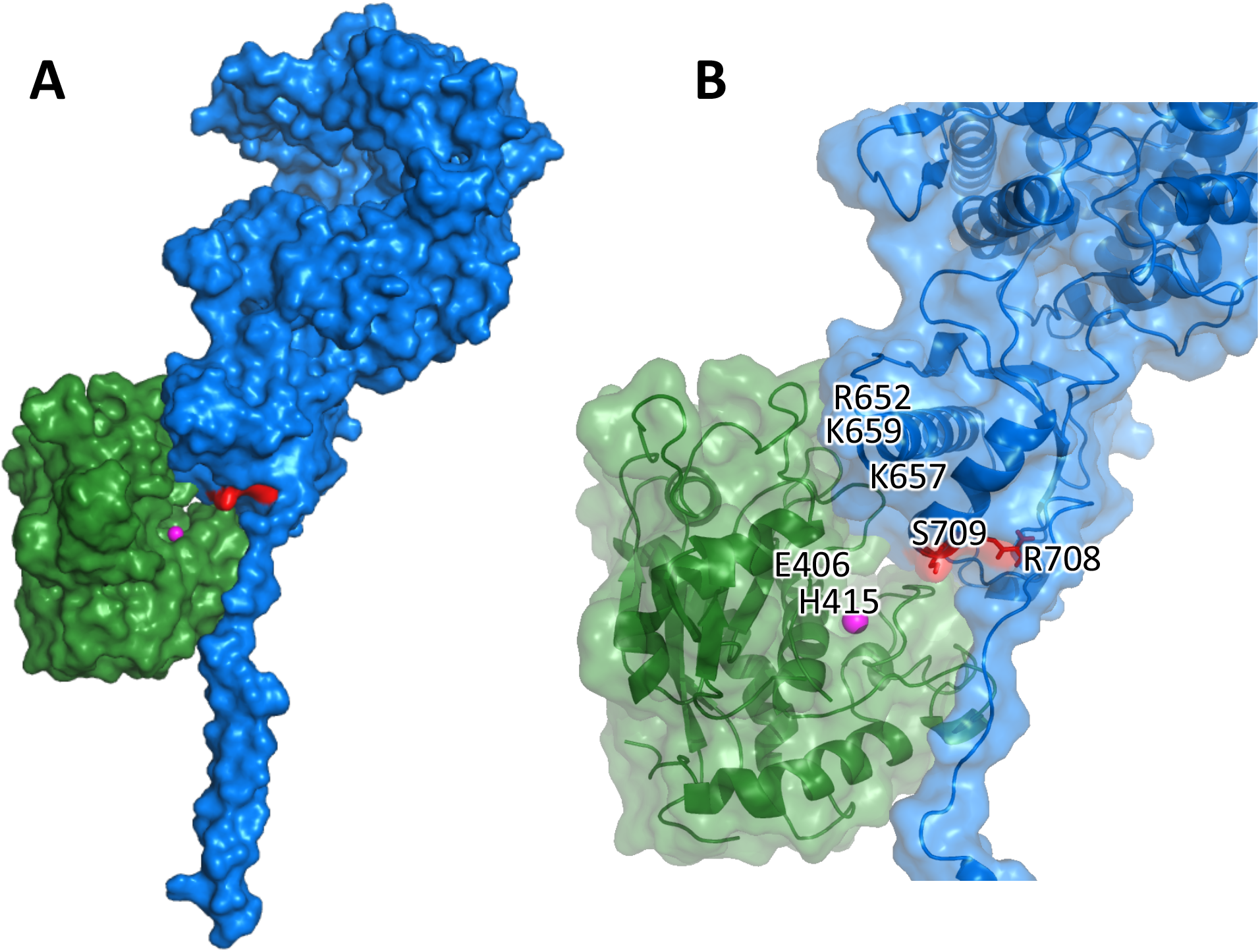
ACE2 monomer target successful docking with ADAM17 ligand. **A**. ADAM17 (green) ligand docking to unobstructed readily accessible monomer ACE2 (blue) neck region residues. **B**. Closeup of ADAM17-ACE2 docking contact interface showing ADAM17 catalytic zinc (magenta) and active site pocket residues H415 and E406 of ADAM17 attractions ≤2-3 Å to ACE2 residues R652, K657 and K659, but >10-15 Å distant to ACE2 R708 and ACE S709 (red). Similar results were obtained with SARS-CoV-2 RBD complexed to the ACE2.

In contrast to monomer ACE2, Fig. 3 shows that ADAM17 active site pocket residues failed to dock appropriately with the dimer-of-heterodimers comprised of two ACE2 subunits complexed with two B^0^AT1 subunits; the results were the same whether the ACE2 subunits were complexed to SARS-CoV-2 RBD (6M18, Fig. 3A,B) or excluding SARS-CoV-2 RBD (6M17, Fig. 3C,D). In either scenario, docking attempts resulted in ADAM17 active site pocket residues exceeding >10-15 Å to ACE2 neck region representative residues ACE2_R652, ACE2_K657, or ACE2_K659.

**Fig. 3.**
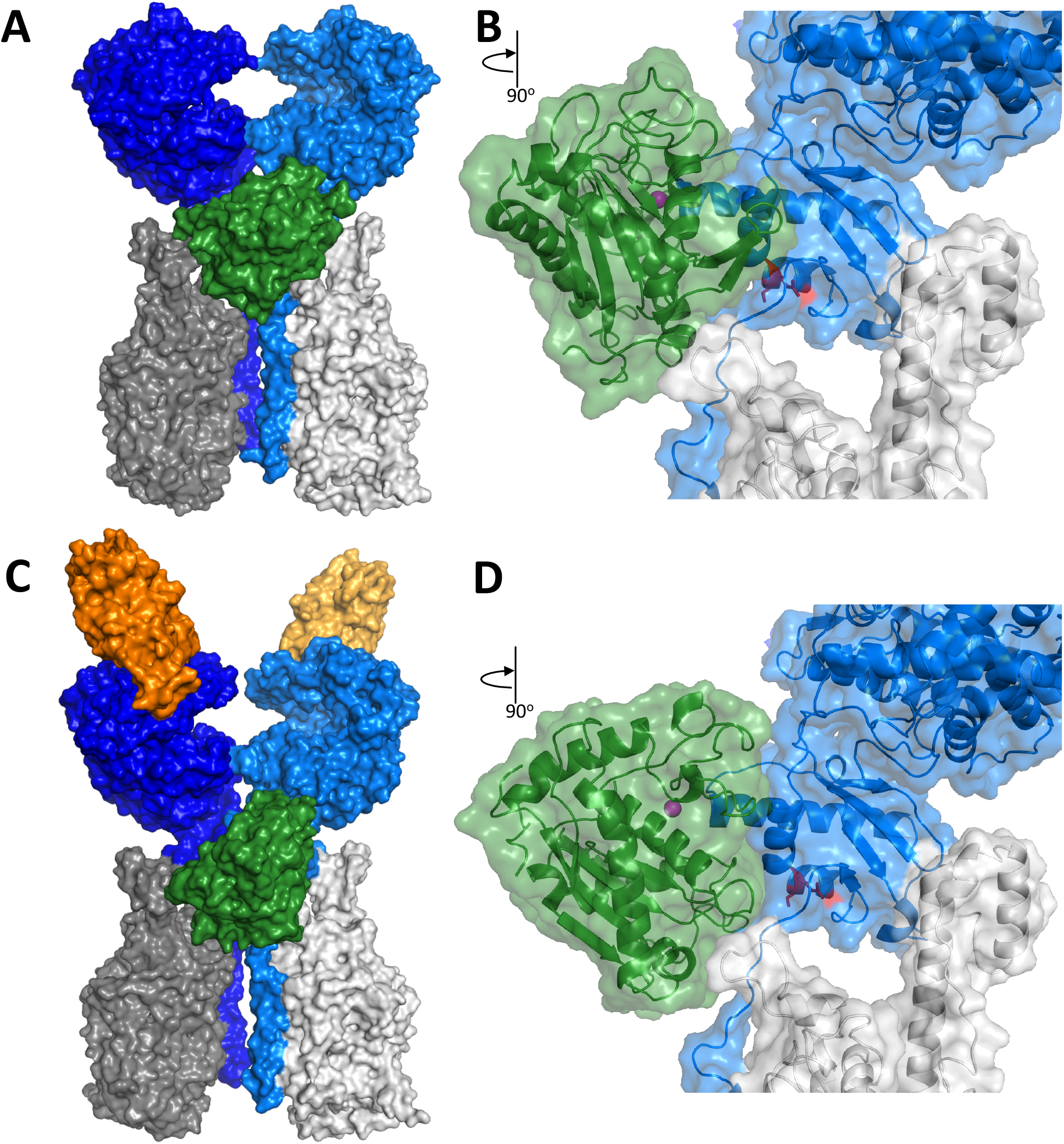
Interference of ADAM17 docking with dimer-of-heterodimeric 2ACE2:2B°AT1 complexes 6M17 or 6M18. **A.** Front view of failed attempted docking contact of ADAM17 (green) ligand with PDB ID: 6M18 dimer-of-heterodimers comprised of two ACE2 subunits (light blue and dark blue) with two B^0^AT1 subunits (light gray and dark gray). **B.** Closeup side view of (A) showing ligand ADAM17 (green) catalytic pocket with zinc atom (magenta) hindered from docking contact (>10-15 Å distance) with ACE2 neck region residues. **C.** Front view of failed attempted docking contact of ADAM17 with PDB ID: 6M17 dimer-of-heterodimers comprised of two B^0^AT1 subunits with two ACE2 subunits complexed with SARS-CoV-2 RBD (gold and brown). **D.** Closeup side view of (C) showing ligand ADAM17 (green) catalytic pocket with zinc atom (magenta) occluded from docking contact (>10-15 Å distance) with ACE2 neck region residues.

To explore the nature of the interferences observed in Fig. 3, ADAM17 ligand dockings were then attempted by employing targets comprised of a single heterodimer comprised of 1ACE2:1B^0^AT1, as shown in Fig. 4. Here, ADAM17 active site residues made successful docking contacts at distances ≤2-3 Å with neck region residues of heterodimer 1ACE2:1B^0^AT1, either without SARS-CoV-2 RBD (Fig. 4A,B) or with RBD (Fig. 4C,D) complexed to the ACE2 subunit, with the exception that ACE2_R708 and ACE2_S709 remained distant. The data of Figs. 3 and 4 were then assembled to reveal the influence of B^0^AT1 on the superposition of ADAM17 docking in relation to ACE2 neck region residues ACE2_R652, ACE_K657, ACE2_K659, R708, and S709 as shown in Fig. 5. The results (Figs. 3A-D and 5A) collectively implicated ADAM17 docking interference as assigned to the presence of the second B^0^AT1 complexed to the neighboring ACE2 of the dimer-of-heterodimers assembly.

**Fig. 4.**
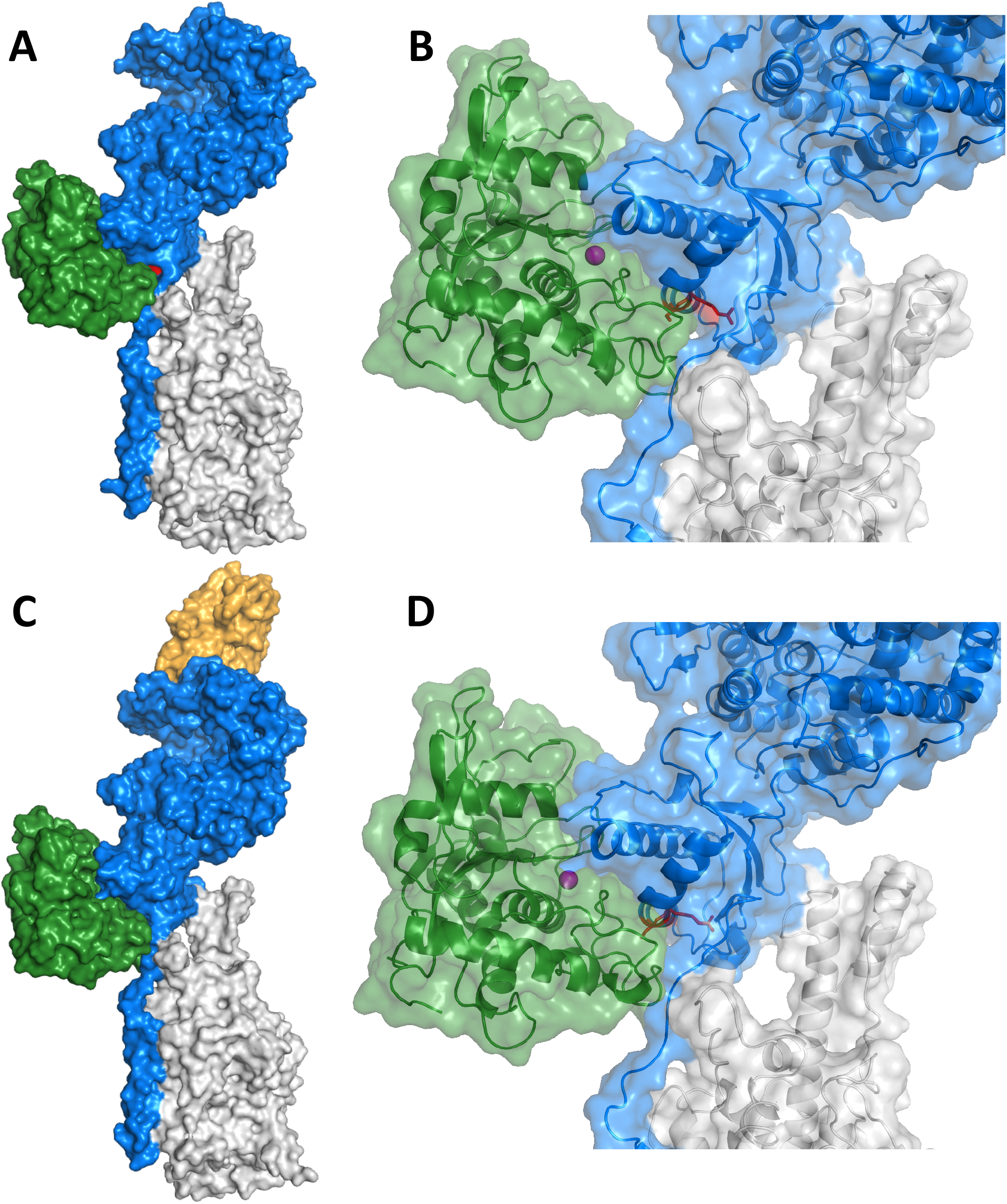
ADAM17 successful docking to 1ACE2:1B°AT1 heterodimer. **A.** ADAM17 (green) ligand successful docking contact with heterodimer of 1ACE2 (blue):1B°AT1 (gray). **B.** Closeup view of (A) showing ADAM17 (green) active site pocket residues (with zinc atom (magenta) docked (<2-3 Å) to 1ACE2:1B°AT1 heterodimer complex. **C.** ADAM17 successful docking to 1ACE2:1B°AT1 complexed with SARS-CoV-2 RBD (gold) similar to (A). **D.** Closeup view of (C) showing docking contact results the same as (B). Although these 1ACE2:1B°AT1 heterodimers do not have physiological antecedence with experimental evidence, these dockings were included for completeness of the combination permutations of ACE2 and B°AT1 subunits.

**Fig. 5.**
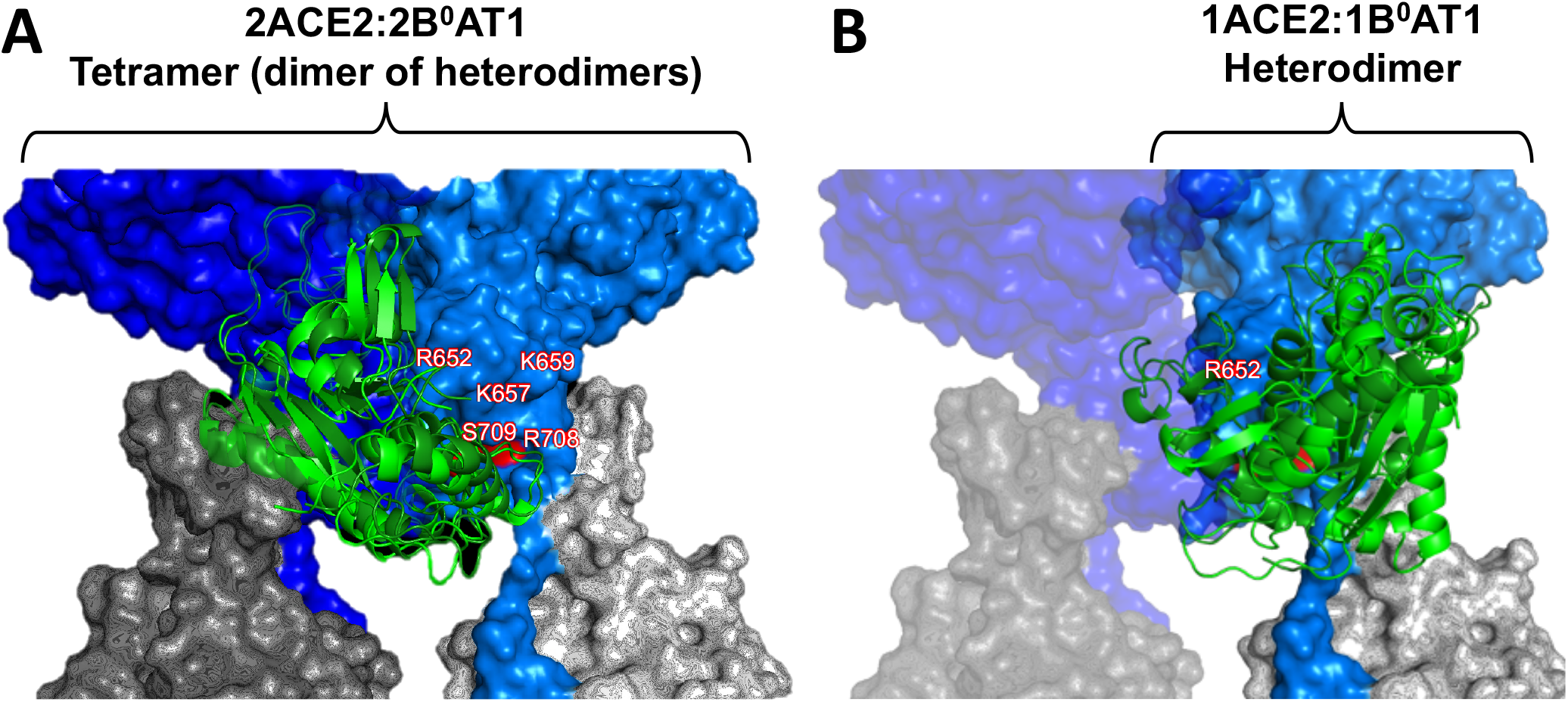
Superposition of ADAM17 docking. **A.** Interference of ADAM17 docking to 2ACE2:2B°AT1 dimer-of-heterodimers (PDB ID:6M18 or 6M17). Presence of neighboring B°AT1 subunit (dark gray at left of A) of partner ACE2 (dark blue at left of A) interfered with ADAM17 (green) successful docking contact with neck residues of the other attached ACE2 (light blue at right of A complexed with partner B°AT1 subunit, light gray at right of A). ACE2 neck residues are representatively shown located at ACE2_R652, ACE_K657, ACE2_K659, ACE2_R708 and S709 each >10 Å distance to ADAM17 active site pocket residues. **B.** Successful docking of ADAM17 to 1ACE2:2B°AT1 heterodimer. Heterodimer components are shown in opaque colors, with transparent ghost of neighboring chains of full dimer-of-heterodimer shown for steric orientation relative to panel (A). Here, ACE2_R652, ACE_K657, ACE2_K659 were ≤3 Å from ADAM17, while ACE2_R708 and ACE2_S709 remained >10 Å distant.

## DISCUSSION

The main findings of this *in silico* molecular docking modeling study are: 1) ADAM17 active site pocket can unimpededly dock and form interface contacts with neck region residues of monomer ACE2, yet the ADAM17 active site remains distant from ACE2_R708 and ACE2_S709; and 2) interference of ADAM17 docking to ACE2 in dimer-of-heterodimer arrangements was directly correlated with the presence of a neighboring B^0^AT1 subunit that is complexed to the partnering ACE2 subunit of the 2 x [ACE2:B^0^AT1] dimer-of-heterodimers—with or without SARS-CoV-2 RBD—representing the expression pattern putatively exclusive to intestinal, renal and cardiomyocyte cell types [1–7]. Experimental evidence is lacking in the literature concerning interactions of ADAM17 with ACE2 in organs that exhibit SARS-CoV-2 tropism and are comprised of these cell types that express B^0^AT1. Although the 1ACE2:1B^0^AT1 heterodimer (note: not the 2 x [ACE2:B^0^AT1] dimer-of-heterodimers) does not have physiological antecedence in the literature, the heterodimer molecular dockings were included in the present project for completeness of the permuted combinations of ACE2 with B^0^AT1 subunits in order to arrive at the ADAM17 superposition results shown in Fig. 5.

*In vitro* experiments [47, 49, 55] have posited that a span covering monomer ACE2 residues R652-I741 may be involved in ADAM17 binding, resulting in the shedding of soluble ACE2 ectodomain away from the cell surface into the extracellular milieu. This was corroborated by the present modeling results that implicated ACE2 candidate residues limited to R652-N718 in its neck region (Fig. 1B). However, the literature is conflicted regarding specific roles of ACE2 neck residues in relation to ADAM17 cleavage or other ADAM17 roles such as neck region competition with alternative ACE2-targeted proteases such as transmembrane serine protease-2 (TMPRSS2). Heurich and coworkers [49] employed a transgenic expression system yielding biochemical experimental data implicating ACE2 residues 652-659 as essential for binding recognition by ADAM17 or TMPRSS2. Although Heurich experiments [49] did not ascertain the exact bonds cleaved for shedding the ACE2 ectodomain, the putative cleavage site was reported to be likely somewhere within its neck span of residues 697-716, specifically excluding R621. Jia et al [55] also utilized biochemical experiments to contend that the putative cleavage could occur in the span of ACE2 residues 716-741. In another set of experiments, Lai [47] implicated cleavage between ACE2 residues R708/S709, with R710 playing a role in presumed binding recognition. The present docking modeling results (Figs. 2–5) indicate that the closest ADAM17 active site residues approached ACE2 R708 was >10 Å for any of the ACE2 conformations tested, whether ACE2 monomer, heterodimer, or dimer-of-heterodimers. Therefore, the present docking modeling is the most consistent with, and primarily supports, the experimental data of Heurich **[49],** and refutes the premise of Lai **[47]** that the cleavage occurs between ACE2 residues R708/S709.

In conclusion, these findings collectively provide the underpinnings and gateway to future wet lab experiments designed to understand the role of B^0^AT1 involvement in COVID-19 and the role of ADAM17 steering ACE2 events. This is especially important relating to the capricious manifestations of COVID-19 in intestinal and renal epithelial cells and cardiomyocytes, with implications useful for consideration in pandemic public hygiene policy and drug development.

